# Stemflow as a Storm-Driven Pathway for Fungal eDNA Transport

**DOI:** 10.64898/2026.10.05.756909

**Authors:** Brian W. Valdman, Brian M. Jusko, David Burke, Sarah Carrino-Kyker, Robert Krebs, John T. Van Stan

## Abstract

Stemflow transports water and biological material from forest canopies to the stem base, yet fungal studies of this flux have largely relied on microscopy of conidia. We used ITS2 metabarcoding to characterize fungal environmental DNA (eDNA) in stemflow from 15 trees – five each of American beech (*Fagus grandifolia*), sugar maple (*Acer saccharum*), and tulip poplar (*Liriodendron tulipifera*) in a closed-canopy forest in northeastern Ohio, USA. Stemflow was sampled during eight rain events in the leafless season (November 2024–May 2025), along with simulated bark and gutter washes to assess near-collar and collector signals. Natural stemflow communities were compositionally distinct from both controls and occupied greater ordination space. Ascomycota dominated across all samples, with *Parafenestella, Cytospora, Microcera, Nectria*, and *Aureobasidium* among the most abundant resolved genera and 13 of the 25 most abundant taxa differed significantly among host species. Woody-plant-associated endophytes, canker fungi, and saprobes dominated ecological guild composition. Storm event explained 28.6% of taxonomic variation, host tree species 24.1%, and their interaction 10.0%. Guild composition was more strongly structured by host species (35.4%) than storm event (20.4%). Lichenized fungi and lichen-associated fungi were detected. Thus, leafless-season stemflow integrates broad bark-associated fungal eDNA, including recurrent lichen-derived material, beyond the morphologically distinctive conidia captured by traditional methods. Stemflow eDNA therefore offers a minimally invasive means of monitoring canopy fungal diversity and the strong host and temporal filtering of DNA exported from woody surfaces.

**Importance:** A rainstorm can turn a tree stem into a fungal sampling device. We show that stemflow integrates fungal eDNA across the woody canopy while preserving signatures of both host identity and storm-to-storm biological turnover, revealing a minimally invasive pathway for repeated monitoring of aboveground fungal communities.

## Introduction

Forest canopies represent interfaces where atmospheric, hydrological and biotic processes converge. Leaves, bark and epiphytic organisms form a porous, biologically active surface that regulates how precipitation, particles, solutes and organisms move from the atmosphere to the forest floor during storm events (Parker 1983; Ponette-González et al. 2020). Within this canopy system, stemflow is a spatially concentrated precipitation flux that integrates material from the contributing crown and can deliver hundreds to thousands of liters per storm to near-stem soil (Sadeghi et al. 2020; Beven and Van Stan 2026). Bark surfaces host abundant and diverse epiphytic plants, cryptogams, and microbiota (Amici et al. 2020; Porada et al. 2023; Barrett et al. 2026) and therefore stemflow processes may annually yield 10^14^–10^16^ bacterial cells (Bittar et al. 2018), 10^5^–10^6^ microfauna (Ptatscheck et al. 2018; Guidone et al. 2021), and 10^9^–10^10^ fungal spores (Magyar et al. 2021) at studied locations. The biological composition of stemflow is theoretically considered the outcome of source availability on canopy surfaces (Bittar et al. 2018), organismal shedding (Teachey et al. 2018), first-flush mobilization (Magyar et al. 2021), particle retention (Van Stan et al. 2021b) and the hydrologic pathways by which water is routed across bark (Mabrouk et al. 2022; Wischmeyer et al. 2024).

Fungi occupy a privileged position in this pathway because they form persistent, surface-bound communities on bark, leaves, lichens, bryophytes, trapped litter, honeydew films and other canopy microhabitats. Some conidia-or ascospore-forming taxa are delivered aerially and captured by wet surfaces, whereas others are members of resident bark and epiphytic communities stabilized by microtopography, biofilms, extracellular polymers and repeated wetting-drying cycles (Ando and Tubaki 1984; Ando 1992; Gönczöl and Révay 2004; Magyar 2008; Sridhar 2009; Chauvet et al. 2016). Lichenized fungi and their lichenicolous associates are especially enduring components of bark communities and form long-lived thalli, host microbial partners, and alter wetting and drying, which can shape stemflow chemistry. The fungal stemflow literature has historically observed the canopy through the lens of rain-mobilized conidia (Magyar et al. 2021), the asexual spores produced on conidiophores, classically by blastic or thallic conidiogenesis (Walther et al. 2005). This conidia-based record is valuable, but it is necessarily selective to flow-competent conidia from exposed microhabitats. Multiradiate taxa such as *Tricladium, Articulospora* and *Tetraploa* are visually distinctive (e.g., see microscopy images from Gönczöl and Révay 2003; Magyar 2008; Sridhar 2009), whereas many elongate scolecosporous forms are difficult to resolve morphologically and are often collapsed into broad categories. More importantly, lichenized Ascomycota, corticioid or yeast-like Basidiomycota, melanized corticolous fungi, endophytes and fragile hyphal fragments may be nearly invisible to morphology-based stemflow sampling even when they dominate bark biomass.

Environmental DNA (eDNA) metabarcoding may change the sampling problem: rather than relying only on recognizable conidia, DNA can be recovered from cells, spores, hyphae, thallus fragments and extracellular material (Beng and Corlett 2020; Sakata et al. 2023; URycki et al. 2024). Stemflow eDNA has already been used to characterize bacterial communities (Teachey et al. 2018), microbial eukaryotes (Gordon et al. 2025b), lichen communities (Sakata et al. 2023) and targeted canopy disease signals (Gordon et al. 2025a), demonstrating that stemflow can carry biologically meaningful information about the surfaces and organisms encountered along its flowpath (Mabrouk et al. 2022). This signal may reflect relatively persistent canopy communities and event-specific inputs. For example, Teachey et al. (2018) found that bacterial communities in net rainfall varied strongly among storm events, with air-mass back-trajectories (Stein et al. 2015) further suggesting that atmospheric source regions influenced the organisms delivered to, and subsequently mobilized from, canopy surfaces. Whether fungal eDNA in stemflow is structured primarily by similarly transient storm inputs or by the more persistent identity of the host tree and its bark habitat remains unknown.

Here we use internal transcribed spacer (ITS2) metabarcoding, taxonomic reannotation and ecological guild analysis (via FungalTraits) to characterize fungal eDNA in stemflow from three co-occurring deciduous tree species. The study addresses several linked gaps. First, morphology-based approaches provide a selective view of stemflow fungi because they emphasize only recognizable, flow-mobilized conidia and may underrepresent bark-resident fungi, lichen-associated lineages, yeasts, hyphal fragments and other poorly resolved material. Second, no studies have examined host-resolved fungal communities across co-occurring tree species with strongly contrasting bark morphologies and stemflow-routing characteristics. Thus, *Fagus grandifolia* Ehrh. (American beech), *Acer saccharum* Marsh. (sugar maple), and *Liriodendron tulipifera* L. (tulip poplar) were selected, as they occupy the same forest stand at Holden Arboretum (Ohio, USA) but represent a continuum from relatively smooth to rough bark. Sampling was restricted to the leafless season (November 2024 to May 2025) to isolate the bark, epiphyte, and bark-associated fungal signals while minimizing contributions from leaf surfaces and foliar fungal communities. Third, simulated near-collar bark washes and collector washes were used to assess whether fungal DNA detected in natural stemflow could be attributed to local bark contamination or collection artifacts. Combined, these methods test (i) how the overall stemflow fungal community is composed; (ii) whether host tree species, storm event, and their interaction structure taxonomic and ecological guild composition; (iii) how lichen-associated fungi and lichenicolous taxa provide evidence of a lichen-derived stemflow signal (Sakata et al. 2023); and (iv) whether natural stemflow carries a fungal eDNA community distinct from that detected in near-collar bark-wash and collector controls.

## Methods and Materials

### Study site and focal host tree species

Storms were sampled from November 2024 through May 2025 at the Holden Arboretum in Kirtland, Ohio, USA, at the Bole Woods plot (41°36′38.8″N, 81°17′27.9″W). The site is a closed-canopy beech-maple forest on somewhat poorly drained Platea silt loam soils with 2–6% slopes (USDA-NRCS, 2025). The regional climate is hot-summer humid continental (Köppen *Dfa*), with a 30-year mean annual temperature of 9.7°C and mean annual precipitation of 1120 mm, distributed evenly across the year (seasonality index = 0.30). Restricting sampling to the leafless season minimized foliar and senescent-litter inputs and focused the eDNA analysis on bark, bark-resident lichens and the dormant-season woody stemflow pathway.

The study used the same 15 mature canopy trees (25–60 cm diameter at breast height) monitored by Noren and Van Stan (2026), comprising five individuals each of *F. grandifolia* (American beech), *A. saccharum* (sugar maple), and *L. tulipifera* (tulip poplar). Bark thickness ranged from beech (2.59%) to maple (5.33%) to tulip poplar (7.47%), a gradient that aligns with bark water storage capacity: low for beech (0.1–1.0 mm; Van Stan et al. 2014), moderate for maple (1–2 mm; Hengst and Dawson 1994), and high for tulip poplar (2.0–3.5 mm; Van Stan and Levia 2010), and with bark conductance, which is expected to vary 40–50% across these species based on relative bark thickness alone (Loram-Lourenço et al. 2022; Ávila-Lovera and Winter 2024).

### Hydrometeorological methods and stemflow sampling

The stemflow collectors used in Noren and Van Stan (2026) were replaced by sterile sampling bins during eDNA collection. Event-scale stemflow was estimated only as hydrologic context, using per-species OLS rain-stemflow (Supplementary Methods S1; Supplementary Table S1). Because source models were calibrated primarily from April–November monitoring and uncertainty bounds reflect slope only, values were treated as order-of-magnitude context for the amount of water and entrained eDNA reaching the stem base. Meteorological data were recorded at 1-minute resolution using a ClimaVue 50 all-in-one weather station (Campbell Scientific, Logan, UT, USA) in an open field approximately 350 m from the study site. The station measured precipitation, air temperature, relative humidity, solar radiation, wind speed and wind direction. Precipitation was measured with the ClimaVue drip-counting gauge at 0.017-mm resolution. Three co-located tipping-bucket gauges (TR-525; Texas Electronics Inc., Dallas, TX, USA; 0.1 mm tip^-1^) provided independent event-scale rainfall measurements. Event totals were compared among gauges to confirm consistency, and the ClimaVue record was used for all analyses.

To identify the synoptic-scale transport pathways and source regions of air masses arriving at the study site, kinematic backward trajectories were calculated using the Hybrid Single-Particle Lagrangian Integrated Trajectory (HYSPLIT) model (Stein et al. 2015) developed by the National Oceanic and Atmospheric Administration Air Resources Laboratory (Supplementary Methods Section S2). The receptor location (trajectory end-point) was defined at the latitude of the site location at Holden Arboretum. The backward trajectories were run for a duration of 48 hours, ending at 00:00 UTC on February 26, 2025. To accurately assess the transport and potential surface deposition of particulate matter (which may contain fungal components), trajectories were initialized at three distinct arrival heights: 500, 1000, and 2000 m above ground level (AGL). These altitudes were selected to capture different vertical layers of the lower atmosphere. The 500 m altitude represents the planetary boundary layer (PBL), where air is well-mixed with the surface and directly dictates local ground-level pollutant concentrations and dry deposition rates. The 1000 m and 2000 m altitudes represent the upper boundary layer and lower free troposphere, which act as primary corridors for the long-range, rapid transport of lofted aerosols (e.g., distant industrial emissions or wildfire smoke).

Each of the 15 study trees was instrumented with a stemflow collection collar constructed from PVC J-channel RV rain gutter drip rail (Tondiamo Inc., China). Collars were attached approximately at breast height (approximately 1.37 m) using 2–4 aluminum nails, spiraled around the full circumference of the bole to produce a continuous gentle slope toward the outlet and sealed above and below with silicone caulk to ensure all water traveling down the stem was intercepted and directed into collections. Stemflow was routed from the collar outlet via a plastic RV gutter spout (Camco Manufacturing, Part 42462, Greensboro, NC, USA) connected to a 2.4 cm diameter silicone tube terminating in a pre-cleaned 10 L high-density polyethylene (HDPE) carboy. Carboys were cleaned between events with a dilute acid rinse followed by repeated rinsing with tap water, dried, and sealed until deployment. To minimize degradation of biological material and reduce the risk of post-collection contamination, carboys were deployed no more than 24–48 hours before a forecasted storm event and retrieved from the field as soon as practicable following the end of rainfall; the maximum elapsed time between storm cessation and sample retrieval across the study period was approximately 40 hours.

### Bark-wash and collar-wash controls

Two potential near-collar contamination sources were identified *a priori*: (1) fungal material eluted from the lower stem surface near the collar installation site, and (2) residual material associated with the collar itself, which despite cleaning between events could harbor residual biological material from prior collections or from ambient deposition during inter-storm intervals. Wash experiments were conducted on separate dry, low-wind days following at least 72 h without measurable rainfall, as verified using the on-site ClimaVue record (Supplementary Figure S1). Four randomly selected study trees per species were sampled during each test (*n* = 12 total trees). Bark washes were conducted on July 29, 2025, and the collar washes on August 6, 2025. Greenhouse well water from the Holden Arboretum was used for both experiments, with a source-water blank collected immediately before each assay. Water was delivered at 1 L min^-1^ (stemflow rates from the study trees maximized around 1.5 L min^-1^ per Noren and Van Stan, 2026) using a peristaltic pump and acid-cleaned sterile silicone tubing. For bark washes, a perforated section of outlet tubing was secured approximately 1 m above the collar, and the intervening bark surface was wetted for 5 min or until 3 L of eluate was collected. For collar washes, water was applied directly to the collection hardware at the same flow rate and target volume. Eluate was collected from the collar outlet in sterile, pre-labeled HDPE containers and processed identically to natural stemflow samples.

### Water filtration, DNA extraction, and ITS2 sequencing

Prior to DNA extraction, each carboy was gently inverted several times to resuspend settled particulates, and 300 mL was withdrawn and processed for eDNA recovery by vacuum filtration through a sterile 47 mm, 0.45 µm mixed-cellulose ester membrane filter (gridded, sterile; Membrane Solutions) mounted in a glass filtration apparatus connected to a vacuum manifold. Following filtration, the membrane was aseptically transferred into a labeled whirl-pak bag, sealed, and placed at -80°C until processing. Between storm sampling events, filtration glassware, forceps, and associated components were decontaminated by immersion in a dilute acid solution (muriatic acid at a 20:1 water-to-acid dilution ratio) for a minimum of 10 minutes, sequentially rinsed with soap, then tap water, and allowed to air dry.

DNA was extracted from filters retrieved from the -80°C freezer, bisected using a sterile scalpel on clean aluminum foil, and each half was transferred into a 1.5-ml bead beating tube (300 mg of 400 µm sterile glass beads; VWR, West Chester, PA, USA, 200 mg of 1 mm sterile glass beads; Chemglass, Vineland, NJ, USA) and CTAB (cetyltrimethylammonium bromide) buffer. Cells were lysed by bead beating (Precellys homogenizer; Bertin Technologies, France) and DNA was extracted using a phenol-chloroform procedure (Burke et al. 2020). Extracted DNA from each sample was suspended in 100 μl Tris EDTA buffer and stored at -80 °C. Extraction controls using chemicals only were run alongside samples.

The ITS2 region was amplified using primers 58A2F (5′-ATCGATGAAGAACGCAG-3′) (Martin and Rygiewicz 2005) and ITS4 (5′-TCCTCCGCTTATTGATATGC-3′) (White et al. 1990) with added Illumina overhang adapters. PCR was performed using Fast Start Taq DNA polymerase (Sigma Aldrich, Saint Louis, MO, USA) at a final concentration mix of 0.5 unit, 0.8mM dNTPs, 0.2 μM of each primer, and 0.5 μg/μl bovine serum albumin in a total reaction volume of 25 μl. Thermocycling included an initial denaturation step of 95°C for 5 min, 25 cycles of 95°C for 30 s for denaturing, 54°C for 60 s for primer annealing, and 72°C for 90 s for elongation, and a final extension of 72°C for 5 min on a S1000 thermal cycler (Bio-Rad Laboratories, Hercules, CA, USA). PCR products were sequenced as 2 x 250-bp reads on an Illumina Mi-Seq V3 sequencer (Illumina Inc., San Diego, CA, USA) through the Case Western Reserve University Genomics Core facility. The sequencing run generated 12.58 million sequence reads in total. Ten samples were removed due to low sequence depth, including two stemflow samples (90 and 106 reads), two bark wash samples (two and 72 reads), and six collar wash samples (between one and 35 reads). The remaining natural stemflow samples had sequence numbers that ranged from 27,299 to 217,833, while reads for the bark wash samples ranged from 10,144 and 213,323 and collar wash samples contained between 895 and 105,584 reads. The remaining 115 samples were processed with the UNOISE pipeline (Edgar 2016b) and sequence reads for these samples are publicly available on the Sequence Read Archive (SRA) of GenBank under BioProject number PRJNA1526776.

Sequences were processed primarily with the USEARCH program (version 11.0.667; Edgar 2010). Paired-end reads were first merged with the fastq_mergepairs command and control PhiX reads were removed with the filter_phiX command. Reads were trimmed of PCR primers using Cutadapt (v4.4; Martin 2011) with up to 2 mismatches allowed during primer removal. Merged, primer-trimmed amplicons shorter than 300 bp in length or with one or more sequence errors were removed with the fastq_filter command, and the orient command set direction by comparing to the UNITE database (version 10.0, release date 2024-04-04; Abarenkov et al. 2024). The unoise3 command created error-corrected and chimera-filtered zero-radius OTUs (zOTUs) where unique sequences with fewer than 8 reads were removed (per the default settings) prior to mapping. The merged reads from each sample with control PhiX and primers removed were then mapped to the zOTUs with the otutab command.

### Taxonomic and Ecological Assignment

Taxonomic assignments of the zOTUs were first made with the SINTAX algorithm (Edgar 2016a) by comparing against the UNITE database. Ecological annotations were then applied to the final zOTU table for natural stemflow samples using FungalTraits as a supporting resource and refined, where appropriate, using taxon-specific literature and lichen resources rather than database labels alone (Nguyen et al. 2016; Polme et al. 2020; Diederich et al. 2025; Consortium of Lichen Herbaria 2026). Lichen-relevant categories distinguished lichenized mycobionts, lichenicolous fungi, possible lichen-associated fungi, melanized corticolous or black-fungal lineages, and unresolved cases. Non-lichen Ascomycota were further summarized into conservative functional groups. The resulting redistribution of reads among host tree species, final taxonomic assignments, and ecological categories is summarized in Supplementary Figure S2.

#### Statistical Analysis

All statistical analyses were conducted in *R* (version 4.5.2; R Core Team 2023). To account for varying sequencing depths among samples, raw sequence read counts were normalized using the estimateSizeFactors function in DESeq2 (version 1.50.2; Love et al. 2014) as recommended by McMurdie and Holmes (2014). Changes in fungal community structure in the stemflow were analyzed with permutational multivariate analysis of variance (PERMANOVA), Permutational Analysis of Multivariate Dispersions (PERMDISP), and principal coordinates analysis (PCoA) in the vegan package (version 2.7.6; Oksanen et al. 2025) of R, all performed on Bray-Curtis dissimilarity matrices calculated with the vegdist function. PERMANOVA was conducted using the adonis2 function with 4999 permutations to assess the effects of tree species, storm event, and back trajectory on the fungal community structure. PERMDISP was conducted using the betadisper function and running permutation tests (function permutest) on the resulting distances to centroid. Changes in community structure were then visualized with PCoA performed with the betadisper function where graphs were constructed using the first two explanatory axes. Covariance ellipses showing the groupings of samples from each tree species were made with the veganCovEllipse function on the PCoA graphs. PCoA graphs were made in ggplot2 (version 4.0.3; Wickham 2016). Multivariate analyses were run on the entire data set (included simulated stemflow samples), as well as on the natural stemflow samples alone with significance considered at ɑ=0.05. The phyloseq package (version 1.54.2; McMurdie and Holmes 2013) was used to make box and whisker plots of observed and ChaoI richness with the plot_richness function and to calculate the relative abundance of the top 20 most common fungal genera by the function transform_sample_counts to calculate relative abundance and then the function glom to agglomerate at the level of genus. A bar graph showing the relative abundance of these genera was made in ggplot2. The relative abundance of the top 20 fungal genera, along with five higher ranked taxonomic groups, for each host tree species were examined with non-parametric Kruskal-Wallis tests where Bonferroni-corrected p-values less than 0.002 were considered statistically significant. Dunn post-hoc tests were conducted for significant Kruskal-Wallis tests with the package FSA (version 0.10.1) and p-values were corrected with the Bonferroni method. The mean and standard error for the relative abundances within each host tree species were calculated with the package doBy (version 4.7.2).

## Results

Stemflow characteristics for eight sampled storm events varied substantially (Table 1a), as did prior climate conditions (Table 1b). Total event rainfall ranged from 4.1 mm to 24.6 mm and wind run ranged from 9.5 to 384.5 km. The highest wind runs corresponded to long storm duration and association with early seasonal changes, and by late-April, wind runs lessened, with the largest rainfall event occurring under notably calm conditions. HYSPLIT back-trajectory analysis revealed that air masses arrived predominantly from southerly directions, with four of eight events classified as having southerly (S) trajectories and the remaining three arriving from the southwest (SW) and one from the northwest (NW). Mean pre-event temperatures ranged from 2.1 to 23.3 °C, RH from 38.6 to 92.3%, and VPD from 0.06 to 1.35 kPa, while winds were strongest during March events and comparatively weak in late April and May.

**Table 1.** Summary of (A) storm event and (B) atmospheric characteristics for the eight storms sampled during the 2024-2025 leafless season at Holden Arboretum, Kirtland, Ohio. Rainfall amount, duration, and mean intensity, as well as wind run were computed from the on-site weather station. Back-trajectories were determined by the NOAA HYSPLIT model (Supplementary Methods S2). Pre-event atmospheric conditions (mean±SD) for the 18-h window preceded rain onset for each sampled storm. Temp. = air temperature; RH = relative humidity; VPD = vapor pressure deficit; Solar rad. = incoming shortwave radiation flux density.

| Event<br>[#] | Date<br>[DD-MM-YYYY] | Amount<br>[mm] | Duration<br>[h] | Intensity<br>[mm h<sup>-1</sup>] | Wind run<br>[km] | Back<br>Trajectory |
| --- | --- | --- | --- | --- | --- | --- |
| 1 | 20-11-2024 | 10.5 | 4.3 | 2.4 | 50.5 | SW |
| 2 | 26-02-2025 | 10.4 | 9.2 | 1.1 | 56.7 | NW |
| 3 | 15-03-2025 | 10.7 | 27.7 | 0.5 | 384.5 | S |
| 4 | 20-03-2025 | 6.5 | 6.2 | 1.0 | 44.8 | SW |
| 5 | 30-03-2025 | 11.9 | 26.2 | 0.5 | 156.4 | S |
| 6 | 19-04-2025 | 4.1 | 8.1 | 0.5 | 48.2 | S |
| 7 | 26-04-2025 | 24.6 | 15.7 | 1.6 | 9.5 | S |
| 8 | 02-05-2025 | 9.9 | 6.8 | 1.5 | 16.3 | SW |

B.
| Event<br>[#] | Temp.<br>[°C] | RH<br>[%] | VPD<br>[kPa] | Wind speed<br>[m s<sup>-1</sup>] | Solar rad.<br>[W m<sup>2</sup>] |
| --- | --- | --- | --- | --- | --- |
| 1 | 12.4 ± 2.4 | 88.3 ± 8.2 | 0.18 ± 0.15 | 1.21 ± 0.76 | 76.2 ± 118.6 |
| 2 | 2.1 ± 1.8 | 92.3 ± 4.9 | 0.06 ± 0.04 | 0.81 ± 0.39 | 53.6 ± 69.5 |
| 3 | 16.6 ± 1.0 | 53.2 ± 5.6 | 0.89 ± 0.15 | 3.67 ± 1.05 | 0.3 ± 1.1 |
| 4 | 18.4 ± 2.9 | 38.6 ± 12.6 | 1.35 ± 0.44 | 2.43 ± 1.39 | 183.4 ± 216.6 |
| 5 | 19.2 ± 0.5 | 58.6 ± 2.0 | 0.92 ± 0.06 | 1.84 ± 0.52 | 49.1 ± 52.0 |
| 6 | 23.3 ± 1.9 | 55.8 ± 13.1 | 1.31 ± 0.50 | 1.69 ± 0.46 | 79.8 ± 164.0 |
| 7 | 20.3 ± 4.6 | 59.4 ± 20.2 | 1.15 ± 0.84 | 0.50 ± 0.65 | 138.5 ± 241.2 |
| 8 | 14.8 ± 6.6 | 67.6 ± 6.5 | 0.62 ± 0.37 | 0.41 ± 0.57 | 178.3 ± 262.8 |

The fungal eDNA community recovered from stemflow was dominated by Ascomycota across all samples, although genus-level composition varied substantially among individual trees and storm events (Figure 1). *Acer saccharum* and *Fagus grandifolia* exhibited particularly heterogeneous communities, whereas *Liriodendron tulipifera* generally showed greater dominance by unresolved Ascomycota. Several identified genera, including P*arafenestella, Cytospora, Microcera, Nectria,* and *Aureobasidium*, reached appreciable relative abundances but varied markedly among samples. Basidiomycota and other non-ascomycete fungi generally represented smaller fractions of the communities, although their abundance occasionally increased in individual samples. Across stemflow samples, 1637 of the 2537 zOTUs (64.5%) and 5.76 million of the 8.98 million reads in the final zOTU table (64.1%) could not be assigned to genus, indicating substantial unresolved taxa. Off-target amplification also recovered algal reads representing some lichen-associated photobiont genera across stemflow samples. PERMANOVA confirmed tree species as a significant predictor of community composition (R^2^ = 0.241, p < 0.001; Table 2).

**Figure 1.**
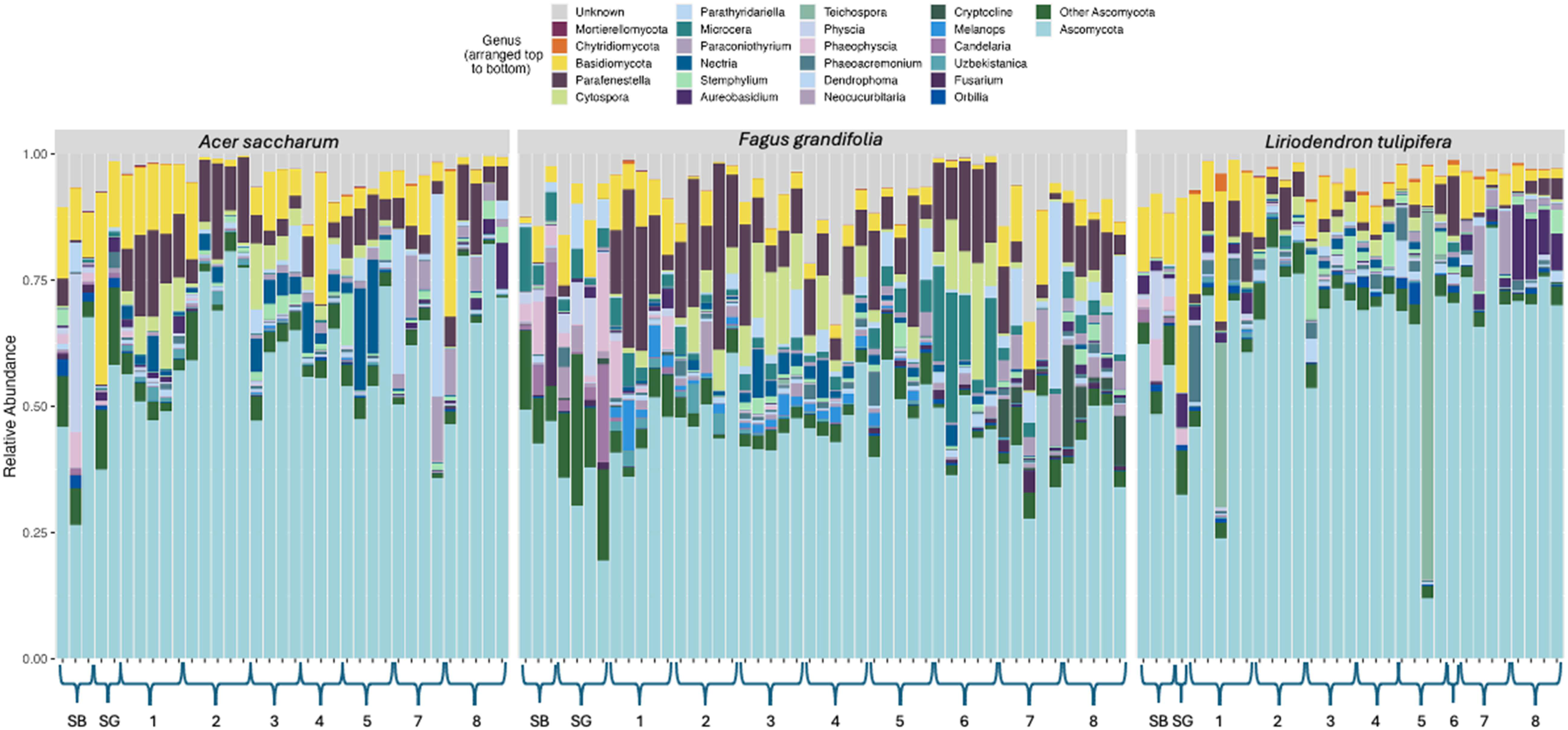
Relative abundance (%) of fungal genera detected in natural stemflow, separated by host tree species. Each bar represents an individual stemflow sample; genera are stacked from top (Unknown) to bottom (*Ascomycota*). Panels reflects both individual tree differences and the chronological progression across storm events (Events 1-8, November 2024-May 2025). “SB” and “SG”: are simulated bark and gutter samples, respectively.

**Table 2.**
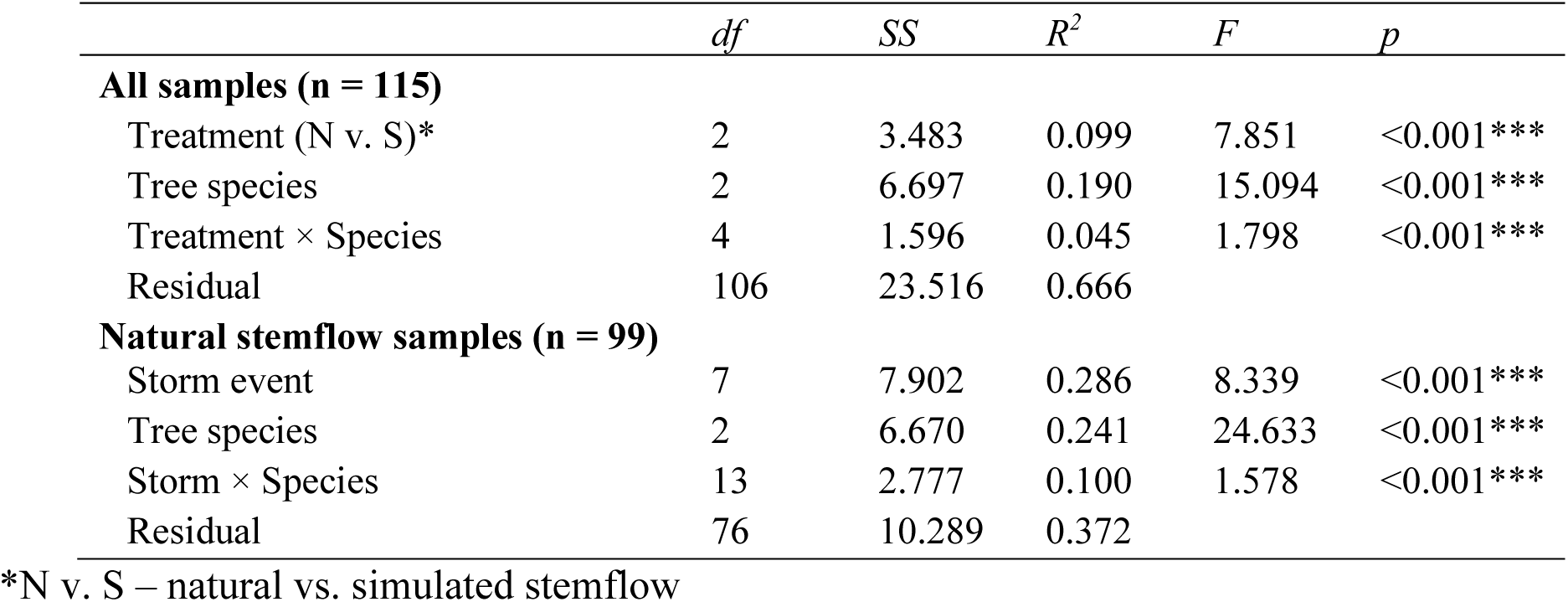
PERMANOVA results for fungal community composition based on Bray–Curtis dissimilarities. The first model included all samples and tested sample type, host tree species, and their interaction. The second model included natural stemflow samples only and tested storm event, host tree species, and their interaction.

|  | *df* | *SS* | *R*<sup>2</sup> | *F* | *p* |
| --- | --- | --- | --- | --- | --- |
| **All samples (n = 115)** |  |  |  |  |  |
| Treatment (N v. S)* | 2 | 3.483 | 0.099 | 7.851 | <0.001*** |
| Tree species | 2 | 6.697 | 0.190 | 15.094 | <0.001*** |
| Treatment × Species | 4 | 1.596 | 0.045 | 1.798 | <0.001*** |
| Residual | 106 | 23.516 | 0.666 |  |  |
| **Natural stemflow samples (n = 99)** |  |  |  |  |  |
| Storm event | 7 | 7.902 | 0.286 | 8.339 | <0.001*** |
| Tree species | 2 | 6.670 | 0.241 | 24.633 | <0.001*** |
| Storm × Species | 13 | 2.777 | 0.100 | 1.578 | <0.001*** |
| Residual | 76 | 10.289 | 0.372 |  |  |

\*N v. S – natural vs. simulated stemflow

Fungal community composition differed among natural stemflow, simulated bark-wash, and simulated gutter-wash samples (Figure 2a). Natural stemflow samples occupied the greatest ordination space and exhibited clear separation among host tree species, whereas both types of simulated washes clustered more closely together. Simulated bark washes were more compositionally diverse than gutter washes (Figure 1; Supplemental Figure S3), but both remained distinct from natural stemflow communities. PERMANOVA confirmed significant effects of sample type, host tree species, and their interaction (Table 2). Sample type explained 9.9% of total compositional variation (pseudo-F = 7.85, p < 0.001), host species explained 19.0% (pseudo-F = 15.09, p < 0.001), and the sample type × species interaction explained an additional 4.5% (pseudo-F = 1.80, p < 0.001). The interaction indicated that compositional differences among natural stemflow and simulated washes varied among host tree species. Dispersion was similar for natural stemflow, simulated bark washes, and simulated collar washes (PERMDISP, p = 0.395).

**Figure 2.**
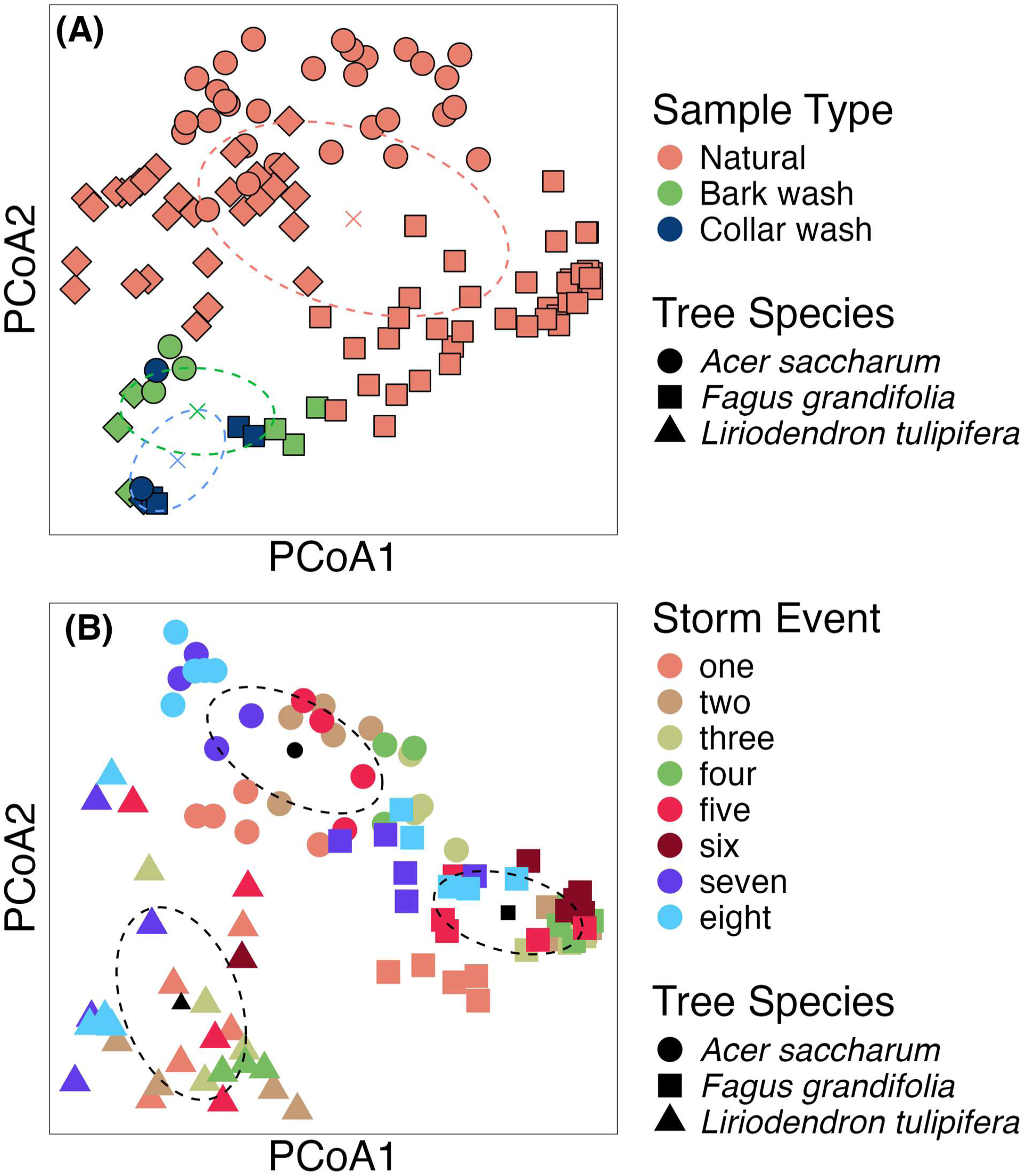
Principal coordinates analysis (PCoA) of fungal community composition based on Bray-Curtis dissimilarities. **(A)** All samples (n = 115), colored by sample type: natural stemflow, simulated bark wash, and simulated gutter wash. Colored centroids and dashed covariance ellipses are shown for sample type. **(B)** Natural stemflow samples only (n = 99), colored by storm event; black centroids and dashed covariance ellipses are shown for host tree species.

Among natural stemflow samples, storm event explained the largest share of variation in fungal community composition (28.6%; pseudo-F = 8.34, p < 0.001), followed by host tree species (24.1%; pseudo-F = 24.63, p < 0.001; Figure 2b; Table 2). The storm event × species interaction explained an additional 10.0% of variation (pseudo-F = 1.58, p < 0.001), indicating that temporal changes in community composition were not uniform among host species. No differences in multivariate dispersion were observed among host species (Pairwise PERMDISP: *F. grandifolia* versus *A. saccharum*, p = 0.31; *F. grandifolia* versus *L. tulipifera*, p = 0.11; *A. saccharum* versus *L. tulipifera*, p = 0.39), indicating that these compositional differences were not attributable to unequal levels of within-group dispersion. There were some differences in multivariate dispersion among storm-event (PERMDISP, p=0.001), most notably a lower within-group dispersion for storm event 6, which was a storm that did not accumulate stemflow from all three tree species (Figure 1). There were also differences in multivariate dispersion between other storm events, indicating that some storms had greater taxonomic variability than others (Figure 2b), although these dispersion differences were idiosyncratic. Broad air-mass back-trajectory class was tested as an alternative representation of between-storm variation. In a model containing host species, trajectory class was significant but explained only 6.2% of community variation (pseudo-F = 4.31, p < 0.001), compared with 25.1% explained by host species. The trajectory × species interaction was not significant (p = 0.133; Supplementary Figure S4). The storm-event model using rainfall explained approximately 4.6 times more variation than the trajectory model (28.6% versus 6.2%) and included a significant interaction with host species. Moreover, trajectory classes were unevenly represented across the seasonal sequence and included only one northwesterly event. Thus, with eight storms, broad trajectory class could not provide a strong explanation of fungal community composition beyond a basic storm-event model.

To identify taxa that differed among host species, Kruskal-Wallis tests were applied to the 20 most abundant fungal genera and the 5 higher-ranked taxonomic bins: 13 of the 25 taxa differed significantly among tree species after Bonferroni correction (Table 3). *Microcera* showed the strongest host association, averaging an abundance of 4.1 ± 0.87% in *F. grandifolia* stemflow but ≤0.22% in the other hosts (χ² = 62.4, p < 0.001). *F. grandifolia* possessed higher relative abundances of *Cytospora*, *Neocucurbitaria*, *Cryptocline*, *Melanops* and *Fusarium. Parafenestella* and *Parathyridariella* were more abundant in *A. saccharum* and *F. grandifolia* compared to stemflow from *L. tulipifera*, whereas Chytridiomycota*, Aureobasidium,* and *Orbilia* were more abundant in *L. tulipifera* stemflow. *Phaeoacremonium* was less abundant in *A. saccharum* than in the other two hosts, and *Neocucurbitaria* was the only taxon for which all three pairwise species comparisons differed significantly. Ascomycota that could not be identified to genus accounted for a smaller proportion of the community in *F. grandifolia* (45 ± 1.1%) than in *A. saccharum* (61 ± 2.2%) or *L. tulipifera* (66 ± 2.9%; χ² = 46.0, p < 0.001).

**Table 3.**
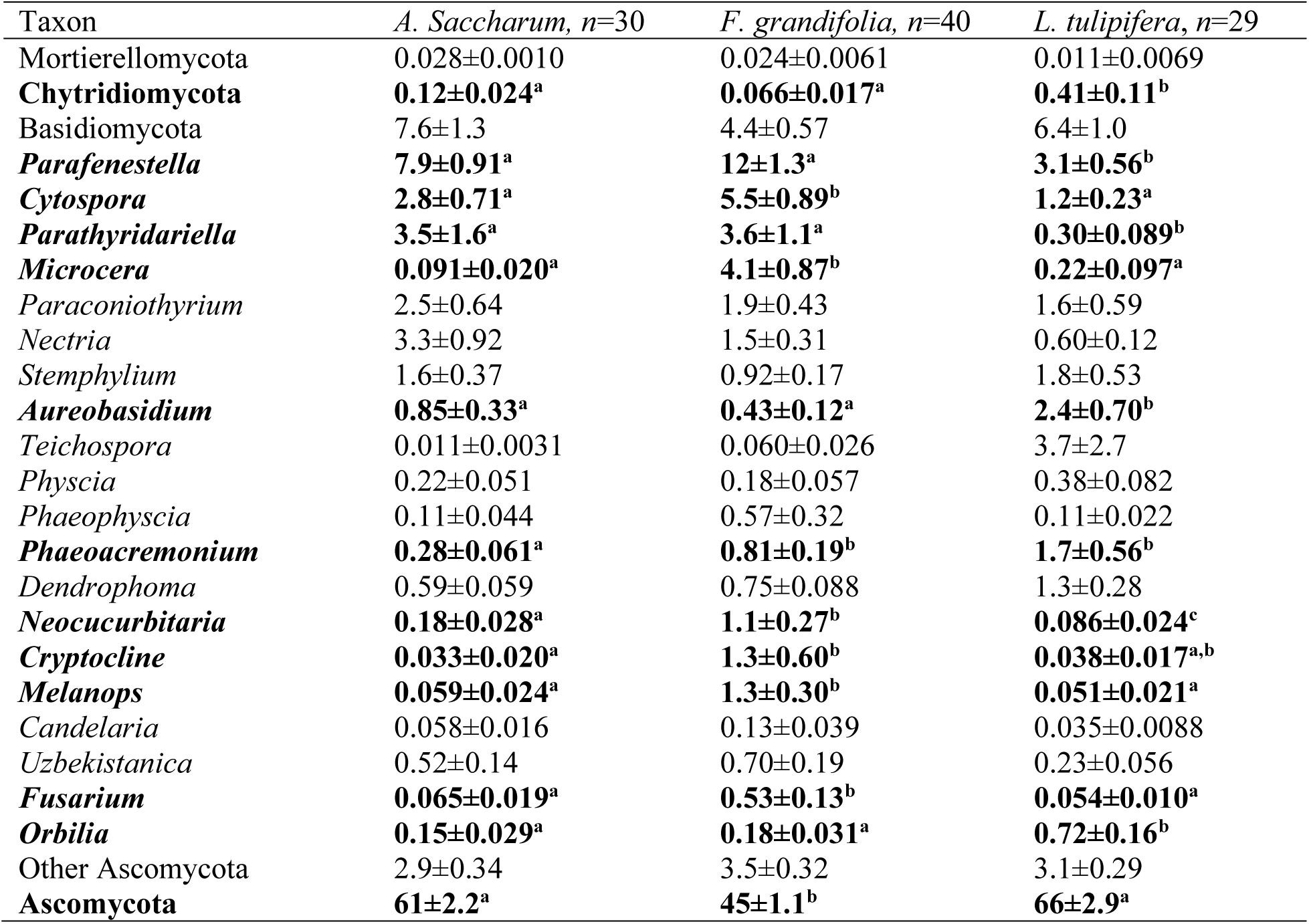
Mean relative abundance (% ± SE) of fungal genera and higher-ranked taxonomic groups in natural stemflow by host tree species. Significant Kruskal-Wallis tests (Bonferroni-corrected *α* = 0.002; 0.05/25 tests) are shown in bold. Where significant, Dunn post-hoc comparisons with Bonferroni-corrected *p*-values are indicated by superscript letters; species sharing the same letter do not differ significantly. “Other Ascomycota” comprises named genera outside the 20 most abundant genera; “Ascomycota” comprises sequences assigned to Ascomycota but not resolved to genus.

Across all curated zOTUs, woody plant canker-, endophyte-, and saprobe-associated taxa comprised the largest fraction of reads in nearly every host-storm combination (Figure 3). Secondary ecological groups varied among host species and events. *L. tulipifera* stemflow possessed a consistently large contribution from wood-, litter-, and soil-associated saprotrophs, particularly during Storms 1–4 and 8. *F. grandifolia* profiles were more variable, with event-specific increases in lichenized or lichen-associated fungi candidates. Stemflow from *A. saccharum* contained comparatively large proportions of fungicolous- or arthropod-associated taxa during several early storms, particularly Storms 2–5, while lichen-related groups increased during Storms 7 and 8. Melanized bark fungi, basidiomycete yeasts and jelly fungi, general environmental taxa, and unresolved sequences generally represented smaller fractions of the total community. The PCoA of sample-level guild profiles showed clear host-associated structure, with the first two axes explaining 53.6% of compositional variation (Figure 4, PCoA1 = 39.1%; PCoA2 = 14.5%). Stemflow samples from *L. tulipifera* occurred predominantly at high PCoA1 scores and low PCoA2 scores, whereas *F. grandifolia* samples were concentrated at low PCoA1 scores. *A. saccharum* samples were more broadly distributed and overlapped with both other hosts. In a corresponding two-factor PERMANOVA, host species explained 35.4% (p = 0.001) of variation in guild composition, while storm event explained 20.4% (p = 0.001), indicating substantial temporal variation superimposed on the stronger host-species pattern.

**Figure 3.**
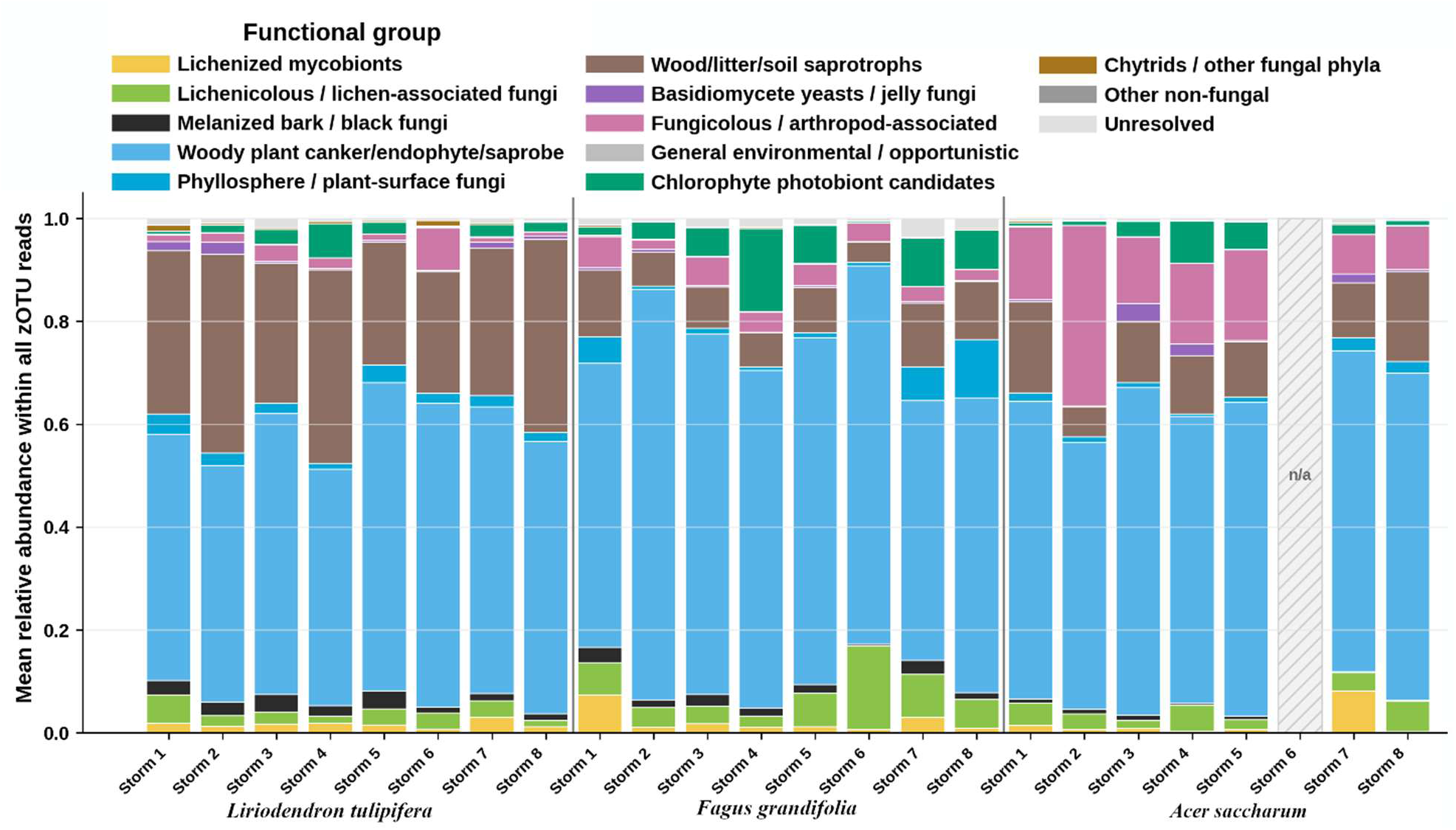
Ecological guild composition of stemflow zOTUs across host species and storm events. Stacked bars show the mean relative abundance of groups within all curated zOTU reads. Chlorophyte photobiont candidates and other non-fungal groups represent off-target ITS2 amplification; these zOTUs were assigned from UNITE taxonomy rather than FungalTraits, which covers fungi only, and are retained here as an indicator of lichen-derived material in stemflow.

**Figure 4.**
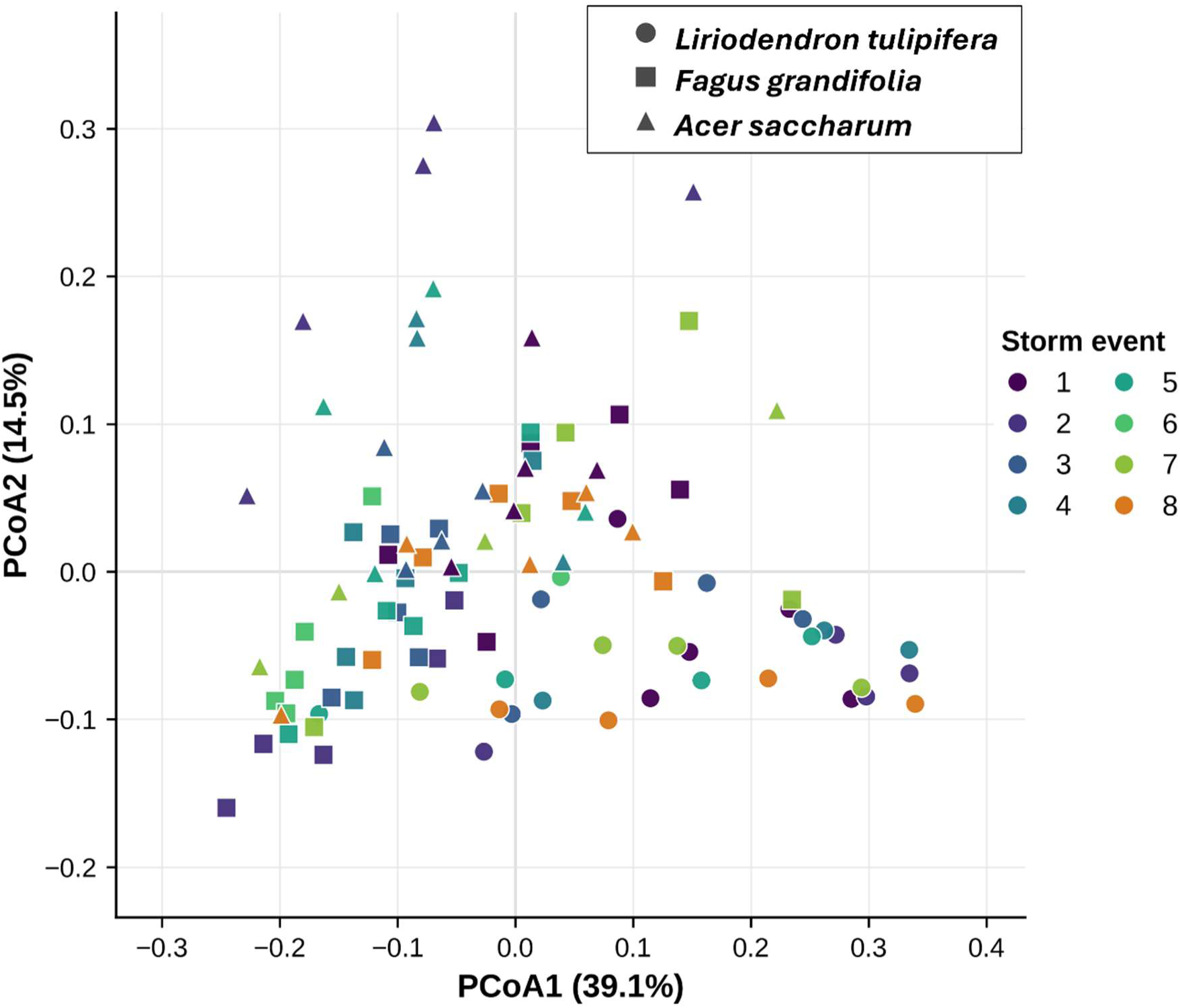
Principal coordinates analysis of ecological guild composition across host species and storm events across all curated taxa explain 53.6% of variation. Samples are colored by storm event. Point shapes denote host species.

## Discussion

Stemflow environmental DNA substantially broadens the fungal information recoverable from canopy drainage waters beyond the morphologically distinctive conidia that have historically dominated stemflow mycology (Chauvet et al. 2016; Ponette-González et al. 2020). Microscopy-based studies have demonstrated that stemflow transports fungal propagules from canopy surfaces (Magyar et al. 2021), but morphology necessarily emphasizes structures that are recognizable and sufficiently intact for identification. Metabarcoding recovered DNA potentially derived from spores, hyphae, thallus fragments, and other fungal materials, extending previous evidence that stemflow can transport bacterial communities (Bittar et al. 2018; Teachey et al. 2018), diverse microbial eukaryotes (Ptatscheck et al. 2018; Guidone et al. 2021; Gordon et al. 2025b), and targeted biological signals of canopy disease (Gordon et al. 2025a), with composition and abundance varying among storms and canopy pathways. That roughly two-thirds of zOTUs and reads could not be resolved to genus is consistent with stemflow being one of the many habitats whose fungi remain poorly represented in reference databases (Hyde et al. 2024), though short ITS2 amplicons and conservative assignment thresholds also limit resolution, so unassigned sequences cannot be equated with undescribed taxa. Together, these fundamental observations support the broader view of stemflow as an integrative conveyor of biological signals and actual dispersal from the canopy to the forest floor (Mabrouk et al. 2022).

Natural stemflow contained fungal communities distinct from simulated bark- and gutter-wash controls, indicating the recovered eDNA signal was not attributable solely to material immediately above the collar or retained on collection hardware. Stemflow eDNA revealed some continuity with classical stemflow microscopy; *Fusarium, Cercospora*, and *Mycocentrospora* had all been recorded from temperate stemflow in Hungary (Gönczöl and Révay 2004). The stemflow community studied here also shared a *Mycocentrospora acerina* observation with tropical Indian stemflow (Sridhar and Karamchand 2009), and tropical mangrove stemflow contained either abundant *M. acerina* in *Avicennia officinalis* L. (Indian mangrove) or *Fusarium-*like conidia in *Rhizophora mucronata* Lam. (red mangrove) (Karamchand et al. 2021). Thus, this northeastern Ohio eDNA assemblage recovered a cross-climate core of rain-mobilized conidial genera. These results highlight the biological diversity transported by stemflow and the extent to which fungal community interpretation depends on database coverage and taxonomic curation. Lichen-associated taxa provide a further example of this broad signal. Lichenized fungi, like *Physcia, Phaeophyscia,* and *Candelaria*, occurred repeatedly across hosts and storms. Corticolous lichens are persistent components of temperate deciduous forest bark communities (Brodo et al. 2002), and bark-surface molecular surveys can recover lichenized fungi that may be overlooked by visual or culture-based methods (Banchi et al. 2018). Bark-associated metabarcoding has likewise demonstrated that molecular sampling can reveal cryptic lichen diversity (Dreyling et al. 2024).

Ecological annotation further indicated that woody-plant canker fungi, endophytes, and saprobes dominated nearly every host–storm combination. Many resolved genera have documented associations with bark or woody plants, including *Parafenestella* (Jaklitsch and Voglmayr 2020)*, Cytospora* (Lin et al. 2024)*, Phaeoacremonium* (Gramaje et al. 2015)*, Neocucurbitaria* (Wanasinghe 2017)*, Melanops* (Phillips and Alves 2009)*, Teichospora* (Jaklitsch et al. 2016)*, Nectria* (Hirooka et al. 2011), and *Aureobasidium* (Untereiner and Malloch 1999; Dreyling et al. 2022). This composition supports long-standing frameworks that bark and canopy microhabitats accumulate fungal propagules between storms that are mobilized by stemflow (Sridhar 2009; Magyar et al. 2016, 2021), while extending recognized diversity to a broader bark-associated fungal assemblage. Critically, many sequences failed to match known species.

The enrichment of *Microcera* in *F. grandifolia* stemflow provides an additional example of the ecological interactions potentially represented by this signal. Members of *Microcera* include insect-associated and lichenicolous fungi (Gräfenhan et al. 2011) and some taxa have been reported from *Physcia* and other lichen thalli (Crous et al. 2021; Cometto et al. 2023). Co-occurrence with *Physcia* therefore raises the possibility that stemflow eDNA can capture interacting members of corticolous communities. Lichenicolous fungi are often cryptic and difficult to identify without specialized examination of individual thalli (Lawrey and Diederich 2003; Muggia and Grube 2018). Molecular sampling of bark-associated material can therefore complement conventional visual and destructive lichen surveys (Dreyling et al. 2024), while stemflow offers the additional advantage of integrating material mobilized over a larger contributing canopy surface area (Sakata et al. 2023).

Host tree species emerged as a major organizer of this diverse fungal signal, explaining 24.1% of taxonomic variation and 35.4% of variation in ecological guild composition. The absence of detectable differences in dispersion among host species further indicates that this pattern reflected differences in community composition rather than greater heterogeneity within one host. Host identity therefore appears to filter both which taxa enter stemflow and the ecological character of the resulting assemblage. Several host properties could contribute to this filtering, although the present design does not separate their individual effects. Bark morphology differs strongly among the three sampled tree species and can alter moisture retention and drying (Rosell et al. 2014; Van Stan et al. 2021a; Ilek et al. 2021), while bark-water relations more broadly affect the microenvironment experienced by epiphytes and microorganisms (Ponette-González 2021; Ávila-Lovera and Winter 2024). Bark pH and nutrient concentrations also vary among tree species (Köhler et al. 2015; Oka et al. 2021), as do porosity and surface structure (Ilek and Kucza 2014; Shtein et al. 2023). These properties influence corticolous lichens distributions and other bark-associated communities (Marmor and Randlane 2007; Westberg et al. 2007; Öztürk and Oran 2011; Lamit et al. 2015; Freudenthal et al. 2024). These habitat differences provide plausible mechanisms for the host-associated variation observed in *Microcera*, *Cytospora*, *Neocucurbitaria*, *Cryptocline*, *Melanops*, *Fusarium*, *Aureobasidium*, and other taxa.

Hydrological routing may provide an additional mechanistic filter. Across the sampled storms, estimated cumulative stemflow was ∼225 L tree^-1^ for *F. grandifolia*, compared with 79 L for *A. saccharum* and 62 L for *L. tulipifera* (Noren and Van Stan 2026). Smooth bark (*F. grandifolia*) can support relatively continuous flowpaths, whereas fissured bark (*L. tulipifera*) or platy bark (*A. saccharum*) partitions water into more spatially restricted pathways and storage zones (Wischmeyer et al. 2024; Katul et al. 2025). Bark-scale flow organization can consequently determine which portions of the woody surfaces contact moving water. In *F. grandifolia*, greater stemflow production may expose a larger or different fraction of the bark-associated community to mobilization, whereas rougher bark may retain water and particulate material within fissures. Bark sorption and retention can further decouple material deposited on woody surfaces from its immediate export in stemflow drainage (Catinon et al. 2009, 2011; Ponette-González 2021). The ecological guild profiles were consistent with such host filtering. *L. tulipifera* contained relatively large contributions from wood-, litter-, and soil-associated saprotrophs, *A. saccharum* showed event-specific increases in fungicolous- and arthropod-associated fungi, and *F. grandifolia* was consistently dominated by woody-plant-associated groups. These categories are ecological interpretations rather than direct demonstrations of activity, but their strong host structure indicates that stemflow is not simply a passive sample of regional airborne spores. Instead, fungal eDNA samples reflect the habitat provided by each tree and the hydrological pathways through which rainwater moves, although these mechanisms remain hypotheses here because bark morphology, hydrological behavior, and host species covary in the present design.

Storm event provided a second, nearly equally strong axis of taxonomic variation. Storm identity explained 28.6% of fungal composition (slightly more than host species), and the significant storm × species interaction explained a further 10%. Thus, fungal communities underwent a common temporal shift across all three hosts where the magnitude or direction of event-to-event change depended partly on host identity. Comparable storm dependence has been reported for bacterial communities transported through forest canopies (Teachey et al. 2018), and temporal structure has also been observed in broader stemflow eukaryotic assemblages (Gordon et al. 2025b). Storm event, however, explained less variation in ecological guild composition (20.4%) than did host species (35.4%), suggesting that taxonomic identities turned over more strongly through time than did the broader ecological character of the communities being mobilized. These storm effects should be interpreted as composite temporal signals rather than responses to rainfall characteristics alone. To focus on bark surfaces, sampling extended from late autumn through spring (leafless season at the site), during which antecedent temperature, atmospheric moisture demand, bark wetness, fungal phenology, atmospheric inoculum, and antecedent dry periods all changed. Seasonal turnover is well-documented in fungal communities associated with temperate forests (Vořiškova et al. 2014), and bark-associated fungi likewise respond to changing environmental conditions through time (Beck et al. 2014). Together, these results suggest fungal eDNA in stemflow records the intersection of two kinds of information: (i) a relatively persistent host-associated signal generated by bark habitat, epiphytes, and hydrological routing, and (ii) a more dynamic event-associated signal reflecting which portions of that community (and potentially which newly deposited organisms) are mobilized during a particular storm. This distinction may explain why host species structured ecological guild composition more strongly, whereas storm identity accounted for slightly more taxonomic variation. Stemflow eDNA therefore expands the observable biological domain of canopy drainage and reveals a temporally dynamic sample of the fungal communities inhabiting and passing through tree surfaces.

## Conclusion

Stemflow provides a hydrologically integrated sample of fungal diversity associated with tree canopies and bark. ITS2 metabarcoding and targeted phylogenetic analysis revealed communities dominated by woody-plant-associated canker fungi, endophyte, and saprobes, alongside lichenized, lichenicolous, melanized, and yeast-like fungi. Host tree species and storm event jointly structured these communities with host identity exerting the strongest influence on ecological guild. Together, these findings extend stemflow mycology beyond morphologically recognizable conidia and establish stemflow eDNA as a minimally invasive means of sampling canopy fungal diversity and its temporal dynamics. Interpretation nevertheless remains constrained by the nature of the analytical methods: ITS2 reads indicate fungal DNA presence but do not distinguish living organisms from dormant spores, dead cells, thallus fragments, or extracellular DNA. Relative read abundance is not equivalent to biomass because copy number, primer affinity, and amplification efficiency vary among taxa. Ecological guilds were inferred from taxonomy and literature rather than measured directly. Because paired bark, lichen, airborne-spore, and throughfall surveys were not done, source contributions cannot be partitioned conclusively. Future studies combining these compartments with quantification and within-storm sampling could test how bark habitat, antecedent drying, and first-flush mobilization regulate fungal export in stemflow.

## Acknowledgements

The authors gratefully acknowledge the support of US NSF DEB-2213623 and the staff at Holden Arboretum.

